## Supplementary figures for "The cryoEM structure of the fibril-forming low-complexity domain of hnRNPA2 reveals distinct differences from pathogenic amyloid and shows how mutation converts it to the pathogenic form"

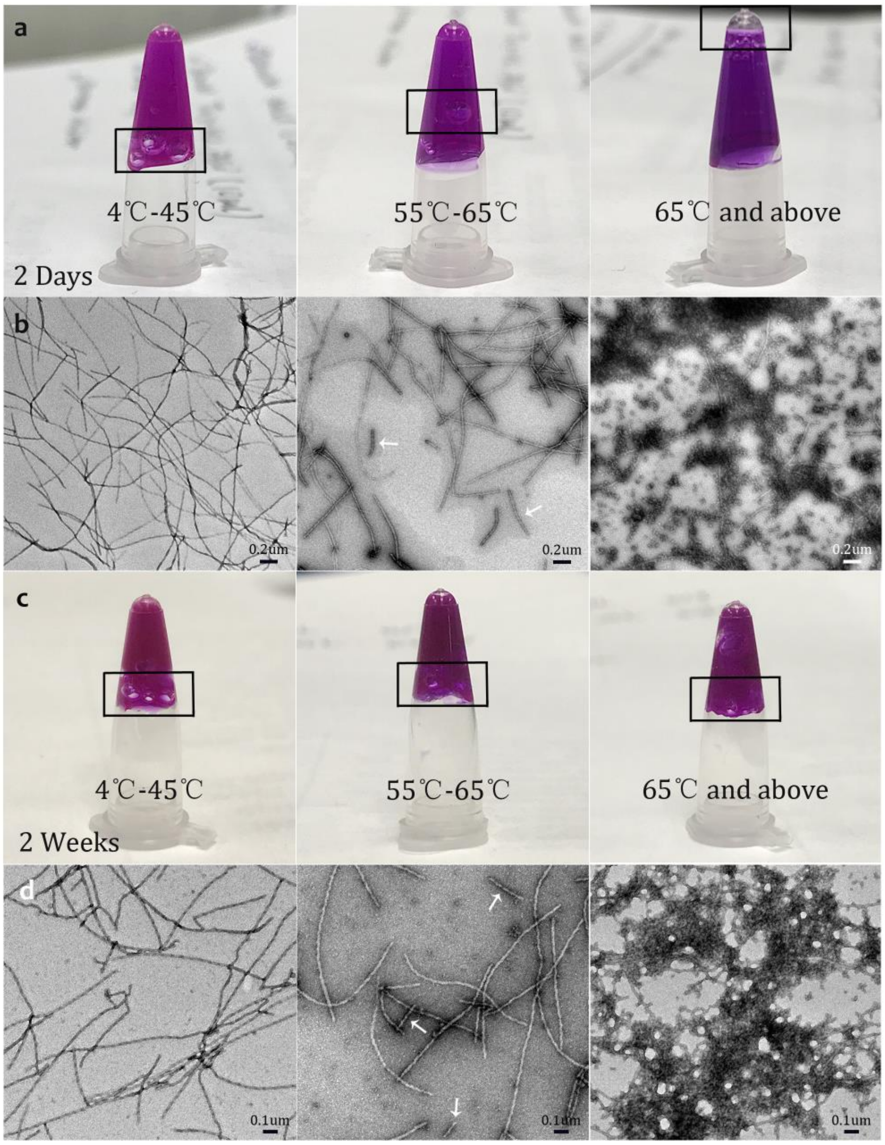


**Supplementary figure1. Demonstration of lability and reversibility of mC-hnRNPA2-LCD fibrils.**

**a.** A hydrogel formed by concentrated mC-hnRNPA2-LCD stays gel-like while heated from 4 to 45℃; it starts to melt at 55℃ as shown by movement of the bubble, and becomes a homogenous solution at 65℃ and above.

**b.** Transmission electron micrographs of 2-day hydrogel droplets at three temperatures: 4-45℃ hydrogel droplets show uniformly similar, amyloid-like fibrils. 55-65℃ hydrogel droplets show fragmented amyloid-like fibrils. 65℃ and above droplets show aggregated fibrils and disk-like structures. Scale bars: 0.2um

**c.** The 2-week hydrogel formed by concentrated mC-hnRNPA2-LCD stays gel-like from 4 to 75℃ and is thus essentially irreversible.

**d.** Transmission electron micrographs of the 2-week hydrogel. Scale bars: 0.1um

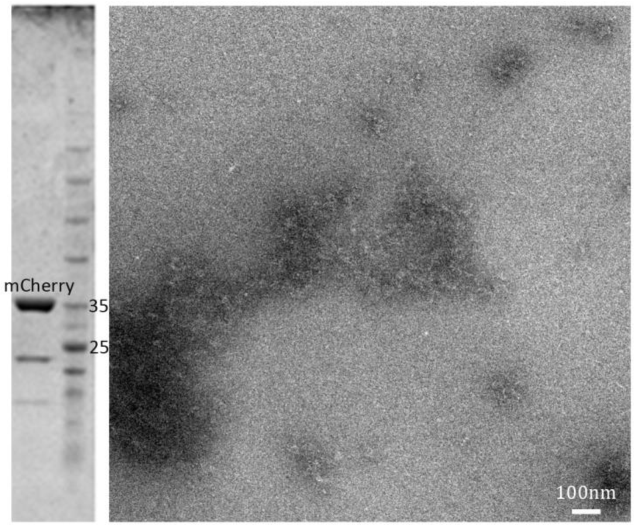


**Supplementary figure2. Negatively stained TEM image of mCherry.**
Left: SDS-PAGE gel with bands showing purified mCherry. Right: Purified mCherry alone heated to 75℃ visualized by transmission electron microscopy shows no fibrils or disk-like structures. Scale bars: 100nm

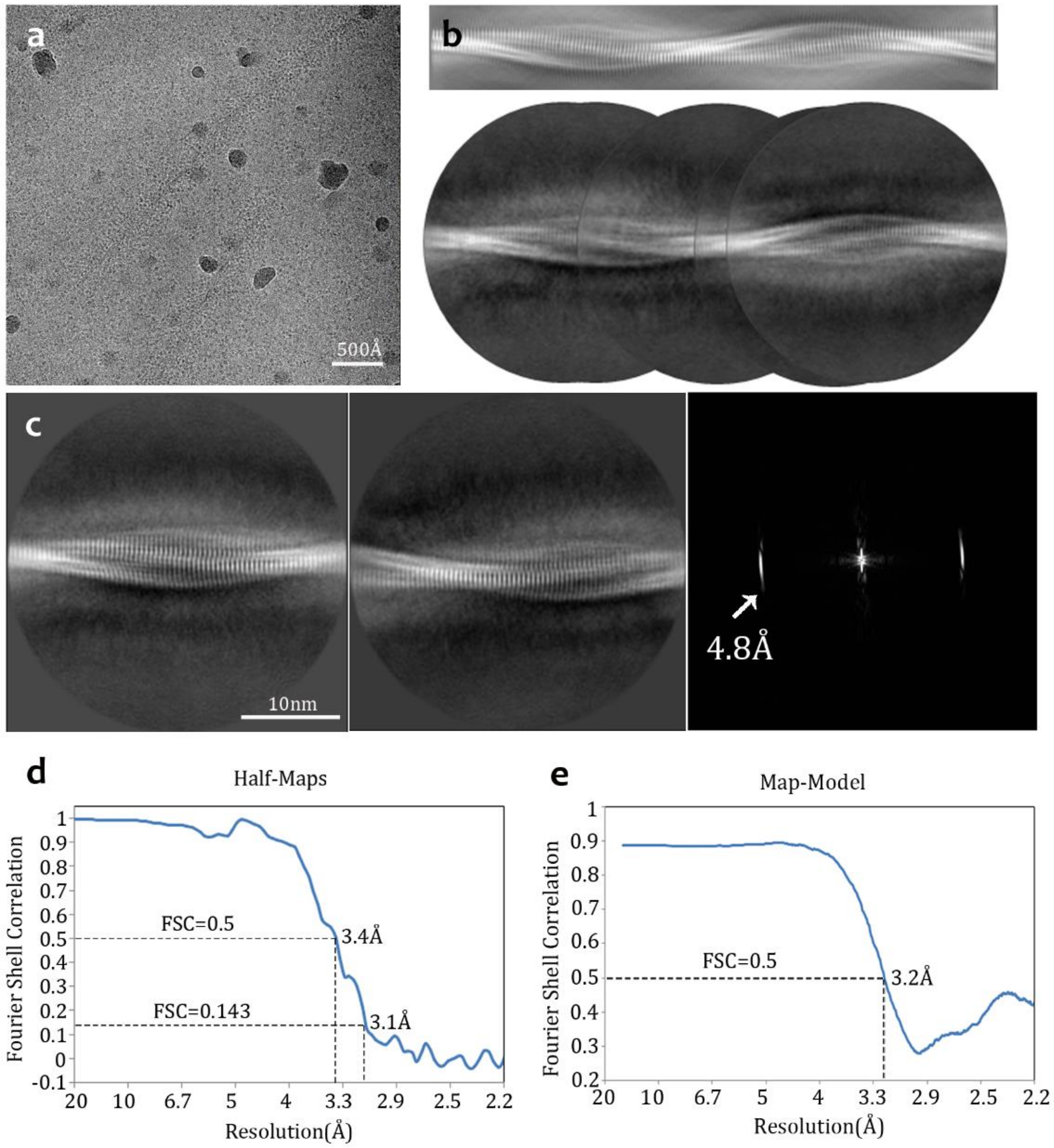


**Supplementary figure3. Cryo-EM data collection, processing, and refinement of mC-hnRNPA2-LCD**

**a.** One representative micrograph from data collection of mC-hnRNPA2-LCD fibrils. Scale bar: 500Å

**b.** Manually assembled full pitch of mC-hnRNPA2-LCD fibril from 2D classification (below) and calculated 2D projection of mC-hnRNPA2-LCD fibril (above).

**c.** Left and middle: two representative images of 2D classifications showing clear 4.8Å layers. Scale bar: 10nm; Right: computed diffraction pattern from 2D class average.

**d.** FSC curve between two half-maps

**e.** FSC curve between the cryoEM reconstruction and the refined atomic model

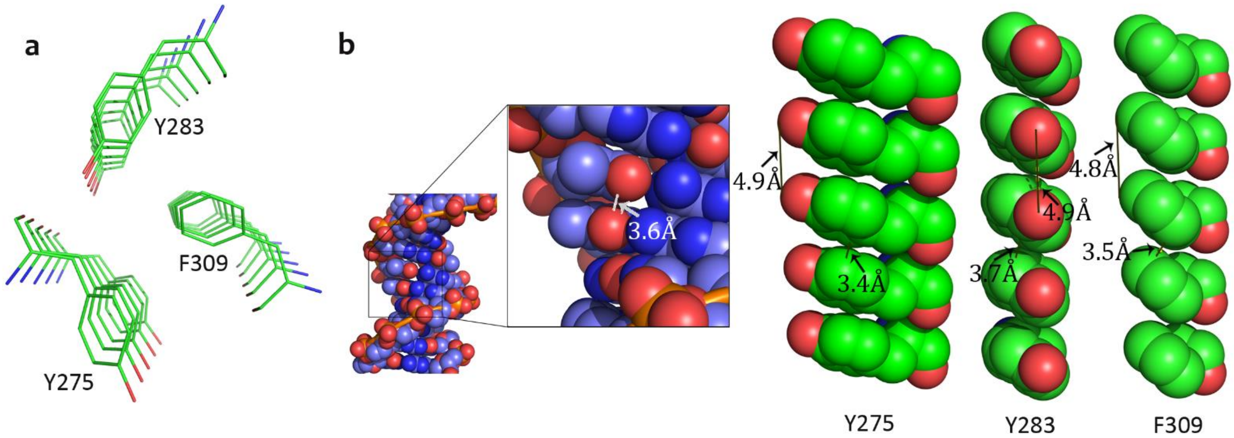


**Supplementary figure4. Detailed interactions of the aromatic triad**

**a.** View down the fibril axis of five layers of the aromatic triad in mC-hnRNPA2-LCD.

**b.** Detailed π-stacking interactions shown in Van Der Waals radii for Tyr275, Tyr283, and Phe309 and a control for B-DNA[67]. Y283 and F309 are having partial Van Der Waals radii contact with distances of 3.7Å and 3.5Å, respectively.

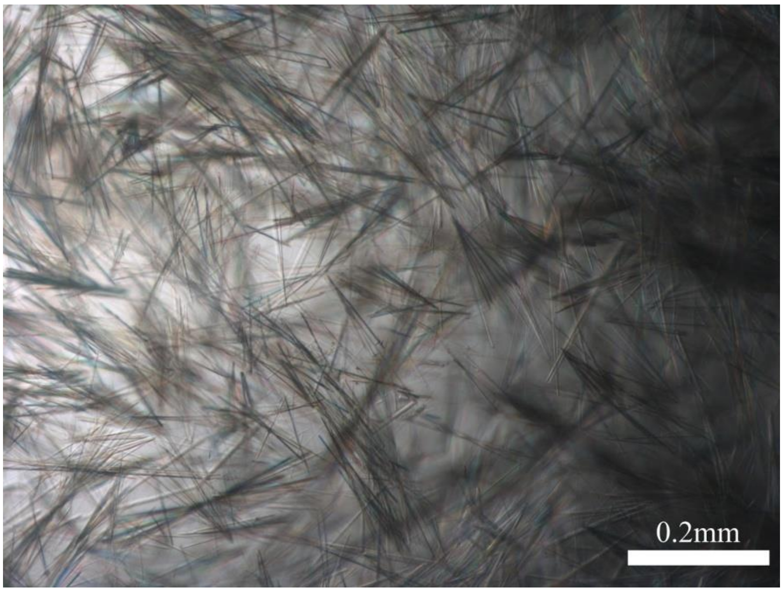


**Supplementary figure5. GNYNVF mutant segment crystals**

Mutant segment GNYNVF needle-like crystals visualized by light microscopy. Crystals growth conditions salt: 0.15M Ammonium Acetate, precipitant: 35% MPD, buffer: 0.1M Bis-Tris, pH 5.5. Scale bar: 0.2mm

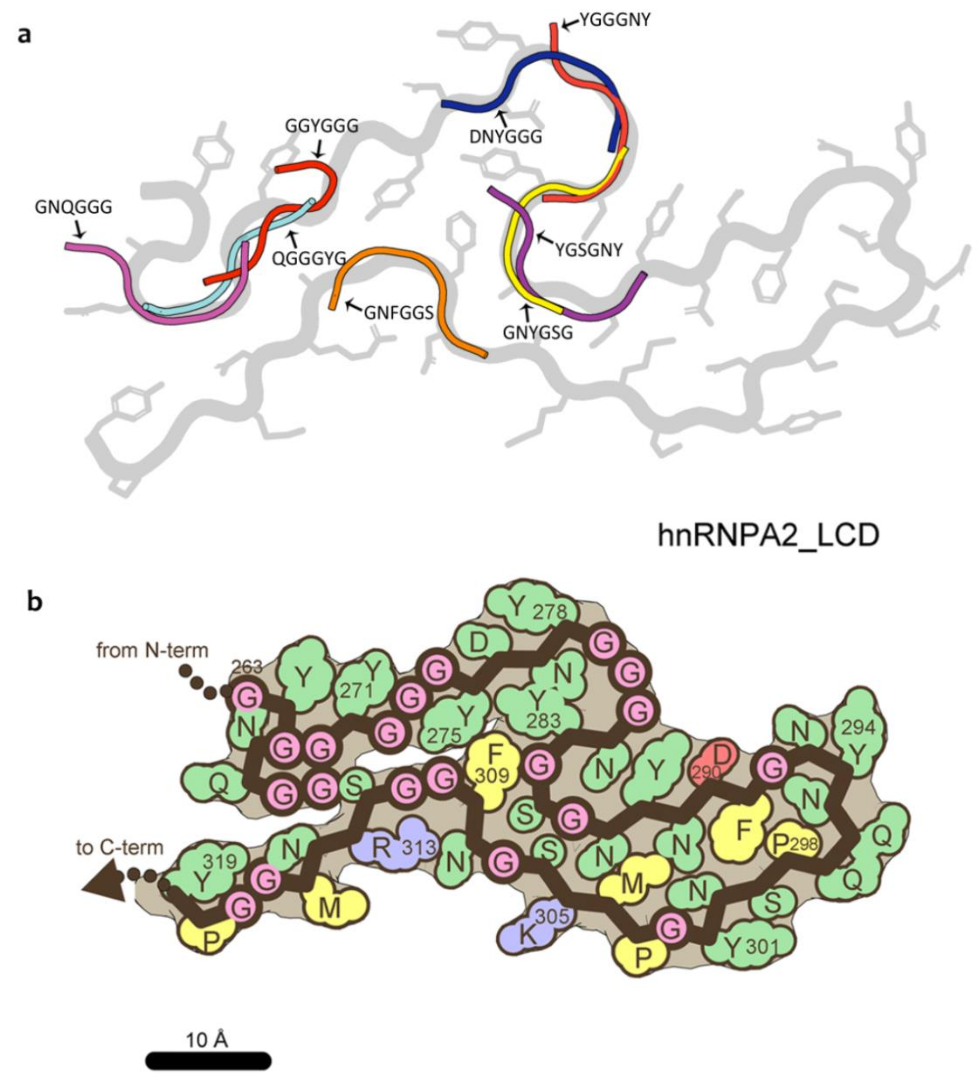


**Supplementary figure6. Structural analyses reveal the basis of mC-hnRNPA2-LCD's reversibility**

**a.** Superimposition of the backbones of 8 predicted LARKS motifs on the atomic model. Arrows show the calculated LARKS structure backbone and the sequences are indicated. mC-hnRNPA2-LCD structure is shown in gray.

**b.** Residue polarity map for the atomic model. Hydrophobic residues colored yellow, polar green, glycine pink, glutamate red and arginine blue.

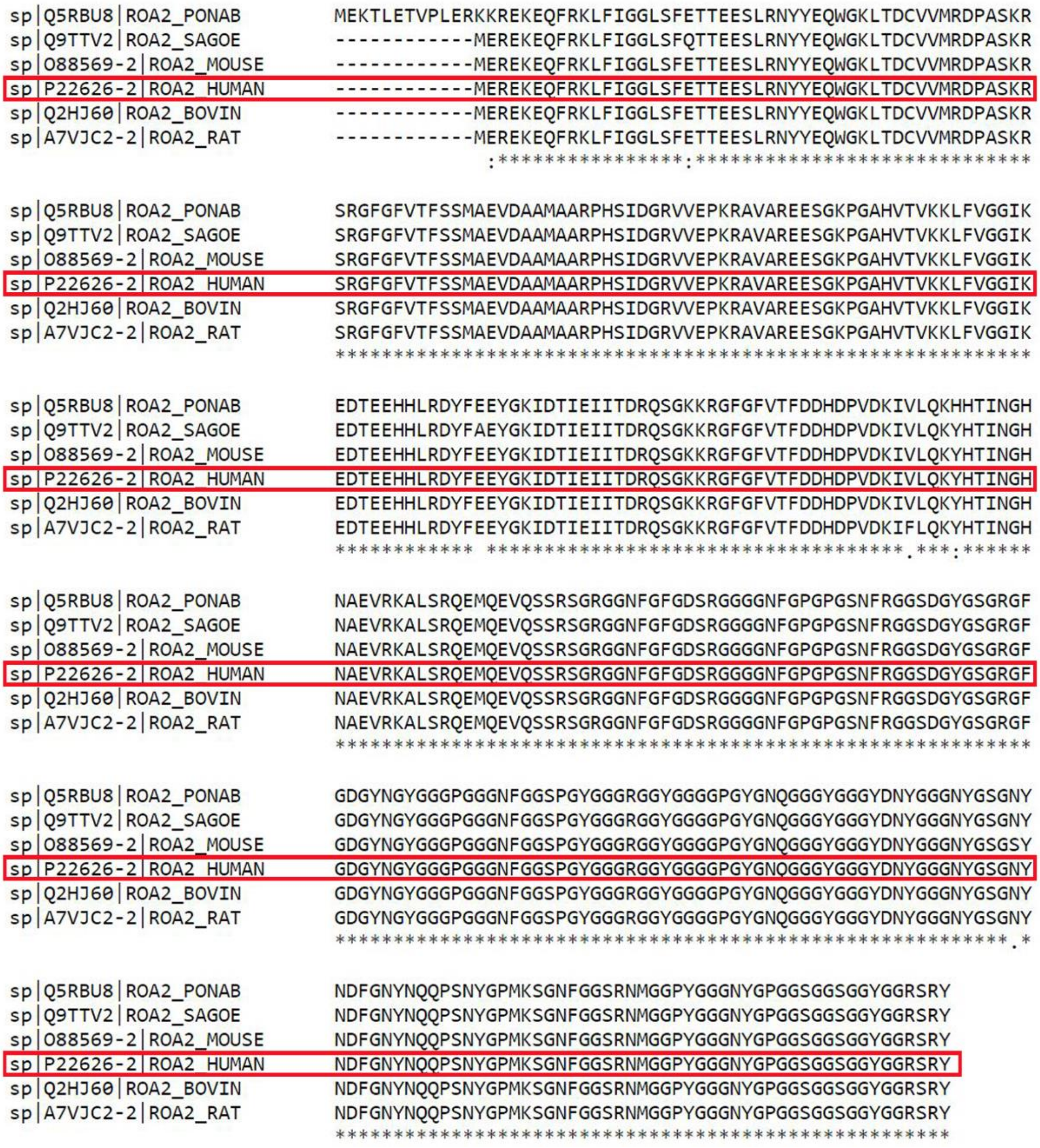


**Supplementary figure7. Sequence alignment of hnRNPA2 of multiple organisms**

Multiple sequence alignment by MUSCLE[68] (https://www.ebi.ac.uk/Tools/msa/muscle/). Human hnRNPA2 sequence is highlighted in red box. PONAB: sumatran oranguta; SAGOE: cotton-top tamarin; BOVIN: bovine. All sequences are from UniProt[69] (https://www.uniprot.org/).

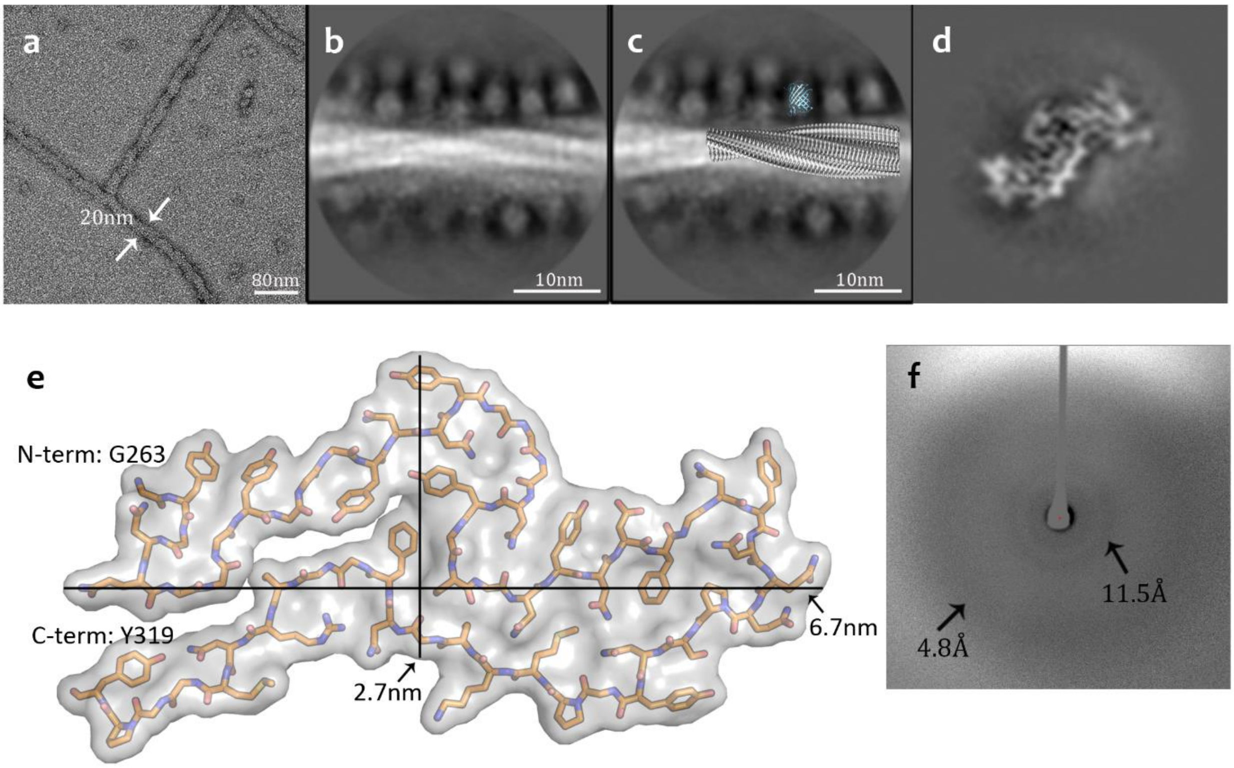


**Supplementary figure8. mCherry forms a fuzzy coat surrounding the hnRNPA2-LCD fibrils**

**a.** mC-hnRNPA2-LCD fibrils visualized by transmission electron microscopy. Scale bar: 80nm

**b.** Representative 2D classification averaged images showing fibrils with a fuzzy coat of spherical blobs.

**c.** Model built with chimeraof the structure of mC-hnRNPA2-LCD (gray) and mCherry (blue) proportional to their real sizes, scaled to B.

**d.** A central slice from the final 3D reconstruction of mC-hnRNPA2-LCD fibril structure.

**e.** Surface model of one cross-sectional layer of the fibril, showing that the length of the model is 6.7nm, and the width is 2.7nm.

**f.** X-ray dififfraction pattern of mCherry solution (hnRNPA2 not included). Two weak reflections are observed at the same Bragg spacings as the mC-hnRNPA2-LCD hydrogel (Figure1), but less sharp. That is, the β-barrel architecture of mCherry produces a diffraction pattern with features that overlap with cross-β diffraction. Attention should be paid when doing X-ray diffraction of tagged proteins if performed in the future.
